# A eDNA-qPCR assay to non-invasively detect the presence of the parasite *Schistocephalus solidus* inside its threespine stickleback host

**DOI:** 10.1101/231670

**Authors:** Chloé Suzanne Berger, Nadia Aubin-Horth

## Abstract

Detecting the presence of a parasite within its host is crucial to the study of host-parasite interactions. The *Schistocephalus solidus* - threespine stickleback pair has been studied extensively to investigate host phenotypic alterations associated with a parasite with a complex life cycle. This cestode is localized inside the stickleback’s abdominal cavity and can be visually detected only once it passes a mass threshold. We present a non-invasive quantitative PCR approach based on detection of environmental DNA from the worm(eDNA),sampled in the fish abdominal cavity. Using this approach on two fish populations (n=151), 98% of fish were correctly assigned to their *S. solidus* infection status. There was a significant correlation between eDNA concentration and total parasitic mass. We also assessed ventilation rates as a complimentary mean to detect infection. Our eDNA detection method gives a reliable presence/absence response and its future use for quantitative assessment is promising.

## INTRODUCTION

Internal parasites with complex life cycles often have multiple effects on their hosts (Marcogliese and Cone, 1997). Studying these parasite-hosts interactions requires to track the parasite during its development in the hosts. One of the model systems for investigating phenotypic alterations of the intermediate host infected by a parasite with a complex life cycle is the *Schistocephalus solidus*-threespine stickleback pair (Barber, 2013). *S. solidus*is a freshwater flatworm that has to infect three distinct hosts to survive and reproduce: a copepod, the threespine stickleback, and a fish-eating bird. The threespine stickleback *Gasterosteus aculeatus* is the specific intermediate host of *S. solidus* and can be infected by one or several worms after consumption of a *S. solidus* infected copepod. Within 24 hours after oral uptake, the small worm penetrates the intestine wall of the fish and migrates into the abdominal cavity (Hammerschmidt and Kurtz, 2007) where it shows a sigmoid growth curve during the first twelve weeks (Barber and Svensson, 2003). When the worm has gained a sufficient mass, it can reproduce in its avian final host (Tierney and Crompton, 1992). Previous studies demonstrated that *S. solidus* infection has major impacts on the stickleback energy demands, physiology, immunity and behaviour (Barber and Scharsack, 2010).

The entire life cycle of this parasite can be reproduced in the laboratory (Barber and Scharsack, 2010). However, the success rate when experimental single infections are used is around 15-20% (Grécias et al., 2016; Weber et al., 2017). Thus, a high number of individuals must be included and assayed in a study to obtain an acceptable final sample size of infected ones. Furthermore, variation in infection rates for sticklebacks caught in the wild is high (Weber et al., 2017), which means that wild individuals considered as uninfected controls may turn out to be infected. A simple, reliable and non-invasive detection method would allow selecting, or avoiding, infected individuals depending on the needs of a specific project. This would allow minimizing the number of individuals used in an experimental procedure, an important aspect for the refinement of animal use protocols.

Available methods to detect sticklebacks parasitized by *S. solidus* relies on morphological modifications that are exacerbated with an increase in parasitic load (Dingemanse et al., 2009). A visual analysis of the level of the abdominal distension of the fish allows to predict the infection status and the parasite index (PI, the proportion of infected fish weight that is contributed by parasite tissue) (Barber, 1997). However, this visual approach is less reliable when the stickleback is infected by a single worm or by a few small worms, which do not induce clear morphological changes. Furthermore, some confounding factors can impact the level of abdominal distension, including eggs and large organs (Barber, 1997). In this context, it is necessary to develop a method that will not be based on morphological criteria.

Our objective was to design a non-lethal method to detect *S. solidus* in the abdominal cavity of its threespine stickleback host. We used two complementary approaches: detection of environmental DNA (eDNA) and measurement of a physiological trait. Our first hypothesis was that *S. solidus* produces cell-free DNA in the fish abdominal cavity, and that this eDNA could be identified by a PCR approach to correctly detect *S. solidus* infection in sticklebacks (Bass et al., 2015). eDNA produced by parasites is a common target in epidemiology to detect infection using a PCR approach in organic samples like serum, feces (Pontes et al., 2002; Xu et al., 2017) or plasma (Imwong et al., 2015). eDNA is also used to detect invasive species in aquatic environments (Takahara et al., 2013). We thus tested a non-invasive quantitative real-time PCR method (qPCR), based on eDNA extraction from fluids of the stickleback abdominal cavity. We designed primers that amplify a gene sequence from the *S. solidus* genome but not stickleback DNA. We tested if we could assign a fish to a given infection status. Since eDNA has been previously used to make quantitative predictions about the biomass of organism detected (Lacoursiere-Roussel et al., 2016), we also tested if eDNA concentration covaried with the total mass of the parasite(s) present within the abdominal cavity of an individual.

Our second hypothesis was that infected fish would show a significantly higher ventilation rate compared to uninfected ones, which could be used to discriminate them. It has been shown that oxygen consumption is higher in infected sticklebacks compared to uninfected ones for a given swimming speed (Lester, 1971) and for fish of similar mass (Meakins and Walkey, 1975). We thus tested the use of the ventilation rate as a rapid and non-invasive indicator of *S. solidus* infection, using a previously published protocol to measure ventilation in stickleback (Di Poi et al., 2016).

## MATERIALS AND METHODS

### Populations of interest

This study was conducted in two populations of threespine sticklebacks. The first population was Lac Témiscouata, Québec (47°40’33”N 68°50’15”O, freshwater environment), where sticklebacks are known to be infected by *S. solidus* and to show typical behaviour alterations (Grecias et al., 2017). The second population was composed of anadromous fish from a salt marsh at Isle-Verte, Québec (48°1’0”N 69°26’59”O; salinity 22-26 ppt (Poulin and FitzGerald, 1989)), for which infection by *S. solidus* is not expected as the worm does not tolerate high levels of salinity (Macat et al., 2015).

### Fish sampling and rearing

Individuals were caught using a seine and minnow traps in Lac Témiscouata as adults in July 2016 and September 2017, and as juveniles in August 2016, Juvenile sticklebacks from the anadromous Isle-Verte salt marsh population were sampled in tide pools in July 2016 using hand nets. Sticklebacks were brought to the “Laboratoire Regional des Sciences Aquatiques” at Université Laval. Adults and juveniles from each population were kept in separate 80 L-tanks under a Light:Dark photoperiod and a temperature of 15°C, reflecting the conditions in their natural environments (Québec, Canada). For anadromous fish, salinity was maintained at 28 ppt. Fish were fed daily with brine shrimps. All sticklebacks were adults at the start of the experiment. They were isolated in 2 L-tanks and were assigned an identification number. 151 sticklebacks were used: 96 from Lac Témiscouata and 55 from Isle-Verte.

### Sampling eDNA from the host abdominal cavity

We sampled the internal fluids of the fish within the intra-peritoneal cavity to obtain eDNA. The fish of interest was taken out of its tank and placed on a sponge. The needle of a syringe filled with 100 μL of PBS (Phosphate Buffered Saline, pH 7.4 Life Technologies) was inserted into its abdominal cavity and PBS was injected. Then, without removing the needle from the abdominal cavity, the plunger of the needle was pulled back until the syringe filled with the PBS that was just injected (up to 100 μL). After the needle was removed, the fish was directly put back in its tank. Between 10 and 100 μL of PBS was obtained and directly added to a tube of 700 μL of Longmire lysis preservation buffer (Longmire et al., 1997). This protocol was done twice for each fish on two consecutive days. On the second day, the PBS used to wash the abdominal cavity was added to the same Longmire tube as the first sample (i.e. up to 200 μL in the 700 μL Longmire tube). Each day, the needle was inserted into a different face of the fish. Abdominal cavity fluid samples were placed at 4°C between the first and the second day, and during the night following the second day. Samples were kept at −20°C before DNA extraction. Fish were fed with brine shrimps after the sampling of the abdominal cavity.

After these two samplings, fish were euthanized with an overdose of MS-222 (75 mg/L mg/kg) followed by exsanguination and dissected to determine if they were infected by *S. solidus*. Fish sex, size and mass, and *S. solidus* mass and number in each fish were noted. The parasite index was calculated using the formula (total weight (mg) of parasites / Total weight (mg) of host plus parasites) x 100. A value of 50 indicates a situation where the total weight of parasites is equal to the net weight of the stickleback (Arme and Owen, 1967).

The protocol was approved by the Comité de Protection des Animaux de l’Université Laval (CPAUL 2014069-3).

### DNA extraction from samples

DNA extraction from the abdominal cavity samples was performed according to an environmental DNA method previously developed (Lacoursière-Roussel et al., under revision), with small modifications to the protocol. To summarize, 30 μL of proteinase K (4 mg/mL) (VWR) was directly added to a Longmire buffer sample and the tube was incubated at 55°C for the night. The sample was transferred into a new 2 mL tube and 950 μL of phenol chloroform isoamyl alcohol (Invitrogen) was added. After shaking and centrifugation, supernatant was transferred into a new tube and 950 μL of chloroform (VWR) was added. After shaking and centrifugation, a maximum of 750 μL of the supernatant was put into a new tube. 750 μL of ice cold isopropanol (Fisher Scientific) and 375 μL of room temperature 5M sodium chloride NaCl were added to the tube, which was placed at −20°C for the night. After centrifugation, all liquid was removed from the tube and 1500 μL of 70% ethanol was added. After centrifugation, ethanol was removed and the tube was air-dried. The DNA pellet was resuspended in 80 μL water and the tube was placed at 55°C for 10 minutes. The tube was placed at 4°C for the night and then stored at −20°C.

### Design of primers specific to the *Schistocephalus solidus* genome

A pair of primers that amplifies a sequence from a nuclear gene (GEEE01010589.1 TSA: Schistocephalus solidus ssol_TR119934_c5_g1_i2 transcribed RNA sequence) from the *S. solidus* genome was designed. The gene sequence was obtained from the de novo transcriptome of *S. solidus* (sequence available on NCBI:https://www.ncbi.nlm.nih.gov/nuccore/GEEE01010589.1?report=fasta) and does not have known homologues in other species (Hébert et al., 2016). The only available information on the function of this gene is that it is highly expressed during early developmental stages of the worm when it is not able to reproduce (Hebert et al., 2017). We targeted the DNA sequence of this gene with the forward for, CGGATTGTCTTCTCGTTGTA and reverse rev-GGACAACCACTGTCCACTAA primers, designed with Primer3 (Untergasser et al., 2012). The primers generated a 200 bp amplicon. We verified with the Basic Local Alignment Search Tool (nucleotide BLAST) on NCBI that the 200 bp amplicon did not align with other sequences except for *S. solidus*.

### PCR assays to determine specificity of amplification

Polymerase Chain Reaction (PCR) assays were conducted on genomic DNA samples to confirm that the designed primers amplified only *S. solidus* DNA and not threespine stickleback DNA. DNA was extracted from *S. solidus* and stickleback samples (Lac Témiscouata) kept at −20°C in ethanol 90%. Tissue samples of 0.3 cm^2^(*S. solidus* body, stickleback tail) were used for the extraction. Extraction was performed using a salt method (Aljanabi and Martinez, 1997). PCR reaction was performed in a total volume of 24.3 μL (13.75 μL of nuclease free water, 1 μL of 10mM dNTPs, 0.25 μL of TAQ DNA polymerase (5U/μL Bio Basic), 2.5 μL of 10x TAQ buffer with MgCL2, 5 μL of DNA (*S. solidus* or stickleback) and 1.8 μL of primer mix (4.56 μM of each primer)). Two negative controls were done (no primers, no DNA). The PCR program was realized with a Mastercycler (Eppendorf) under these conditions: a denaturation step of 2 min at 95°C, an annealing and elongation step of 40 cycles including 20 sec at 95°C, 40 sec at 56°C and 60 sec at 72°C, and a final step of 10 min at 72°C. PCR products were visualized on a 2% agarose gel using SYBR safe staining (Invitrogen).

### qPCR assays

eDNA extracted from the abdominal cavity samples of fish was amplified with a real-time quantitative polymerase chain reaction (qPCR) using the designed primers specific to *S. solidus*. Each 96-well plate included eDNA samples extracted from the body cavity and a standard curve obtained from a pool of *S. solidus* genomic DNA (3 individuals) diluted at different concentrations. The concentrations of the 7-point standard curve ranged from 0.9 ng/μL to 0.01 ng/μL with a dilution factor of 2. qPCR reaction was performed in a qPCR mix including 12.5 μL of PowerUp SYBR Green Master mix (Life Technologies) and 1 μL of primer mix (4.56 μM of each primer). For the standard curve, genomic DNA samples, 6.5 μL of RNAse free water and 5 μL of DNA were added to the qPCR mix. For the eDNA samples, no water was used and 11.5 μL of solution containing the resuspended eDNA was added to the qPCR mix (for a total qPCR mix volume of 25 μL in each case). Two negative controls (no primers, no DNA) were done in each plate. Each reaction was done in triplicate. qPCR was performed with a 7500 Real-Time PCR system (Life Technologies). The amplification was realized according to the conditions described by (Lacoursière-Roussel et al., 2016): 2 min at 50 °C, 10 min at 95 °C, followed by 70 cycles of 15 sec at 95 °C and 60 sec at 56 °C. A melt curve protocol was then performed with temperature going from 60 celsius to 95 celsius to detect primer dimers, check specificity by determining that a single amplicon was produced, and obtain a Tm value for that sample, which can be compared to the standard curve of *Schistocephalus* DNA.

### Ventilation rate measurements

A subset of individuals used in the *S. solidus* eDNA detection experiment was used to quantify ventilation rates (captured in Lac Témiscouata in July 2016, n=20). Ventilation measurements were repeated three days in a row with individuals sampled in the same order each day. For each individual fish, a 250 mL beaker was filled with 50 mL of clean water from the supply used to fill experimental 2 L-tanks. The individual was filmed from the side (JVC Everio camera) to be able to detect opercular movement. At the end of the three days of measurements, these fish were euthanized and their infection status determined. Dissection showed that 13 individuals were infected and 7 were parasite-free. All the infected fish were also detected as infected by the eDNA approach (see below). Films were subsequently analyzed at 0.46 X speed by an observer (NAH) blind to treatment using the open source VLC Media Player (Videolan). The first 30 seconds of the movie where the fish was clearly visible were used to measure ventilation rates. If the individual moved in a position where it was not visible during the first-30-second sampling period, the ventilation measurement was made for the usable seconds within that period and reported back to 30 seconds. The observer repeated all measurements for the first day of sampling to assess accuracy. Measures done twice on the same fish on the same video were highly correlated (Spearman r = 0.99) with most samples having the exact same number of ventilations when measured twice, with a maximum difference of 3 ventilation movements and an average difference of 0.7. A single measurement per video was thus used. A ventilation rate was calculated by dividing the number of ventilation measured in 30 seconds by 30. This measurement was used for further statistical analyses. The total number of ventilation measured during a 30 second interval was used to estimate the number of ventilation over 1 minute for graphical purposes and for comparison to previous studies.

### Analyses

Following qPCR amplification, the presence or the absence of *S. solidus* eDNA in the body cavity samples was determined by comparing the Ct values (amplification curves) and the Tm values (melt curves) of the samples with the ones of the standard curves. A Ct (“cycle threshold”) is the cycle at which a sample reaches a preset threshold of fluorescence, which is the proxy for the quantity of eDNA. A sample with a low Ct means that there were more templates in the original sample and that it thus reached the threshold sooner in the 70 PCR cycles. A body cavity sample showing at least one of its Ct values (among the triplicates) above 0 in combination with the associated Tm value(s) close (+/- 2) to the mean Tm of the standard curve (79.64) was considered positive for *S. solidus* eDNA. On the contrary, if a Ct value above 0 was associated with a Tm value more distant to the mean Tm of the standard curve, then this value was considered negative for *S. solidus* eDNA and the result of primer dimer formation. Finally, if the body cavity sample had only “undetermined” Ct values, it was considered negative for *S. solidus* eDNA, irrespectively of Tm values. For each fish, qPCR results were analyzed with regard to its infection status (parasitized or not by *S. solidus* upon dissection). A true positive was obtained when there was concordance between the infection status and the qPCR results. A false negative was defined when the fish was infected by *S.-solidus*, but no traces of *S. solidus* eDNA were found using qPCR. A false positive was associated with fish not infected by *S. solidus* but for which *S. solidus* eDNA was inferred using qPCR results. All individual results are found in supplementary data.

Statistical analysis was done in R version 3.3.3 (R Core Team, 2017) using R Studio (Version 1.0.136) and the ggplot2 package (Wickham, 2009). To determine if our method allows a quantitative estimate of parasite mass, we tested if the eDNA concentration obtained for infected fish (using the Ct value) negatively correlated with their total parasite mass, using a non-parametric Spearman correlation (a small Ct value should be associated with a larger parasite mass and vice-versa if the eDNA method is quantitative). We used the lowest Ct measured for a given fish if more than one triplicate amplified DNA. Repeatability of ventilation rates over the three days was calculated following a previously developed method (Lessells and Boag, 1987). We compared ventilation rates between infected and non-infected individuals using a t-test, as they followed a normal distribution (based on a Shapiro-wilk test). We tested if ventilation rates covaried positively with the parasite load (parasite index) using a non-parametric correlation (Spearman correlation).

## RESULTS AND DISCUSSION

In parasite-host system studies, being able to detect a parasite during its development in the host is important to better understand their interactions. Here, we demonstrate the power of a method based on detecting eDNA from an internal parasite directly sampled from the abdominal cavity of its specific intermediate host.

### Primers specificity

PCR tests performed with primers that target a sequence from the *S. solidus* nuclear gene NCBI ID GEEE01010589.1 confirmed the amplification of *S. solidus* genomic DNA, and not of threespine stickleback genomic DNA. The PCR product was sequenced using Sanger ABI3730xl (Centre Hospital Universitaire du Québec) to confirm the amplification of the 200 pb amplicon in the *S. solidus* genome. Controls (no template, no primers) were negative as expected.

### Detection of *S. solidus* eDNA in the abdominal cavity of infected fish

qPCR was performed in combination with a eDNA extraction protocol to try to detect *S. solidus* presence in the stickleback abdominal cavity. Standard curves realized with a pool of *S. solidus* genomic DNA samples diluted at different concentrations had mean Ct values ranging from 29.65 (highest concentration) to 36.86 (lowest concentration) and a mean Tm value of 79.64. All the controls were negative. The earliest Ct value we obtained for abdominal cavity eDNA samples with our protocol and amount of starting material varied between 34.4 and 49.6 for positively identified individuals. Using this approach on two fish populations, 148 fish out of 151 were correctly assigned to their infection status (infected or not by *S. solidus*, 98 % of true positive).

In the Témiscouata population, 35 fish out of 96 were infected by *S. solidus.* Single and multiple infections (up to 6 worms) were reported upon dissection, with a median of 1.5 worm, illustrating the predominance of single or double infections. Note that one fish harboured 12 worms, but we did not obtain reliable mass for the worms and as such is included in the dataset only to calculate the infection status assignation rate. Among the infected sticklebacks, 18 exhibited a single worm upon dissection and no external visual clues suggested the presence of *S. solidus* in the fish body cavity (Figure 1). For all infected fish, the total parasite mass was comprised between 130 mg and 2356 mg (median of 495 mg) and the parasite index was always below 50 (between 4.9 and 40.4, median = 16,6%), meaning that the total weight of parasites was always below the net weight of the fish host. 32 out of the 35 infected fish were successfully detected as infected using qPCR. For the 3 remaining infected fish, no trace of *S. solidus* eDNA was detected with our protocol (1.98% of samples tested, low false negative rate). They had no eDNA detected in any of the triplicates (Ct “undetermined”). The 3 fish had a total parasite mass that span almost the whole range found in the study (168 mg and 274 mg for the single infections and 1644 mg for the double infection, from the second smallest to the third largest). Their parasite index (respectively 12.5, 6.5 and 25.3) was above the minimal values from infected fish that were correctly detected. These observations indicate that these false negatives are not due to a lack of sensitivity of the method at low parasitic weight. We rather propose that the critical step of our method is during the sampling of the internal fluids of the fish intraperitoneal cavity. *S. solidus* is free in the abdominal cavity of its host. It is possible that the PBS injection was done too far from the worm to allow the sampling of its DNA. To overcome this issue, we performed body cavity sampling on two consecutive days, and each time the needle was inserted into a different face of the fish. This approach was quite reliable and probably resulted in our low false negative rate. However, we suggest that the eDNA sampling protocol could be extended over three days in order to maximize *S. solidus* DNA capture.

**Figure1.**
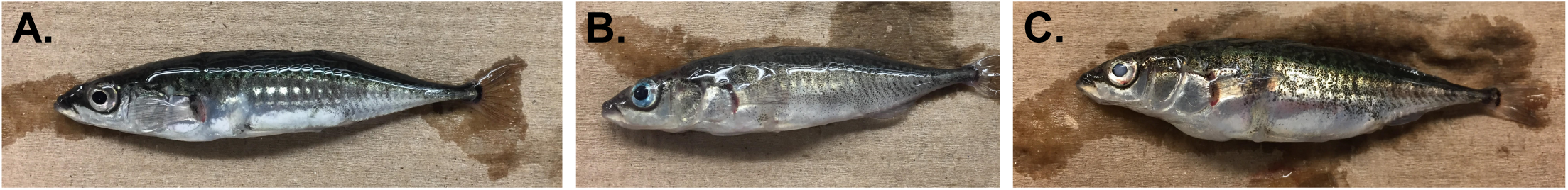
External morphology of threespine sticklebacks from Lac Témiscouata that were infected by *S. solidus* (single or multiple infections) or not. A. Non-infected fish. **B.** Infected fish with 1 *S. solidus* worm (parasite mass 618.2 mgh PI 13.2). **C.** Infected fish with 5 *S. solidus* worms (total parasite mass 2356 mgh PI 39.3). A and B do not exhibit abdominal distension while it is clearly visible in C. Fish B and C were both detected as infected using the eDNA method.

None of the fish from the anadromous population were infected by *S. solidus* (n=55). We never detected *S. solidus* DNA in the samples of the uninfected fish from this population, nor from the 61 uninfected fish from Lac Témiscouata (0 false positive). False positive could have resulted from a lack of specificity of the primers used. The absence of false positive for 116 samples confirms the specificity of the method. A false positive could also have resulted from a signal of an exposed stickleback, i.e. a fish that was exposed to a *S. solidus*-infected copepod, but for which infection did not succeed and the worm did not develop in its abdominal cavity. The absence of false positive suggests that our method does not allow to detect sticklebacks that have been exposed but that did not develop an infection. However, all the fish used were of wild origin, which prevents us to have a record of exposure over the life of an individual fish. Testing this possibility with laboratory-exposed fish could add to the usefulness of the present detection method.

We found a significant correlation between Ct values (which are inversely correlated to the amount of *S. solidus* eDNA) and the total parasite mass found within a fish (one-sided Spearman correlation, rho= −0.41, p = 0.01, Figure 2). This result suggests that in the future this approach could be used in a quantitative way to estimate not only the presence but the total mass of parasites within a host. The correlation is non-linear and there is large variation in total parasite mass for a given Ct value and thus further refinement would be needed. Therefore, our method in its current state gives a very reliable presence/absence response and its use for quantitative assessment looks promising for the future.

**Figure 2.**
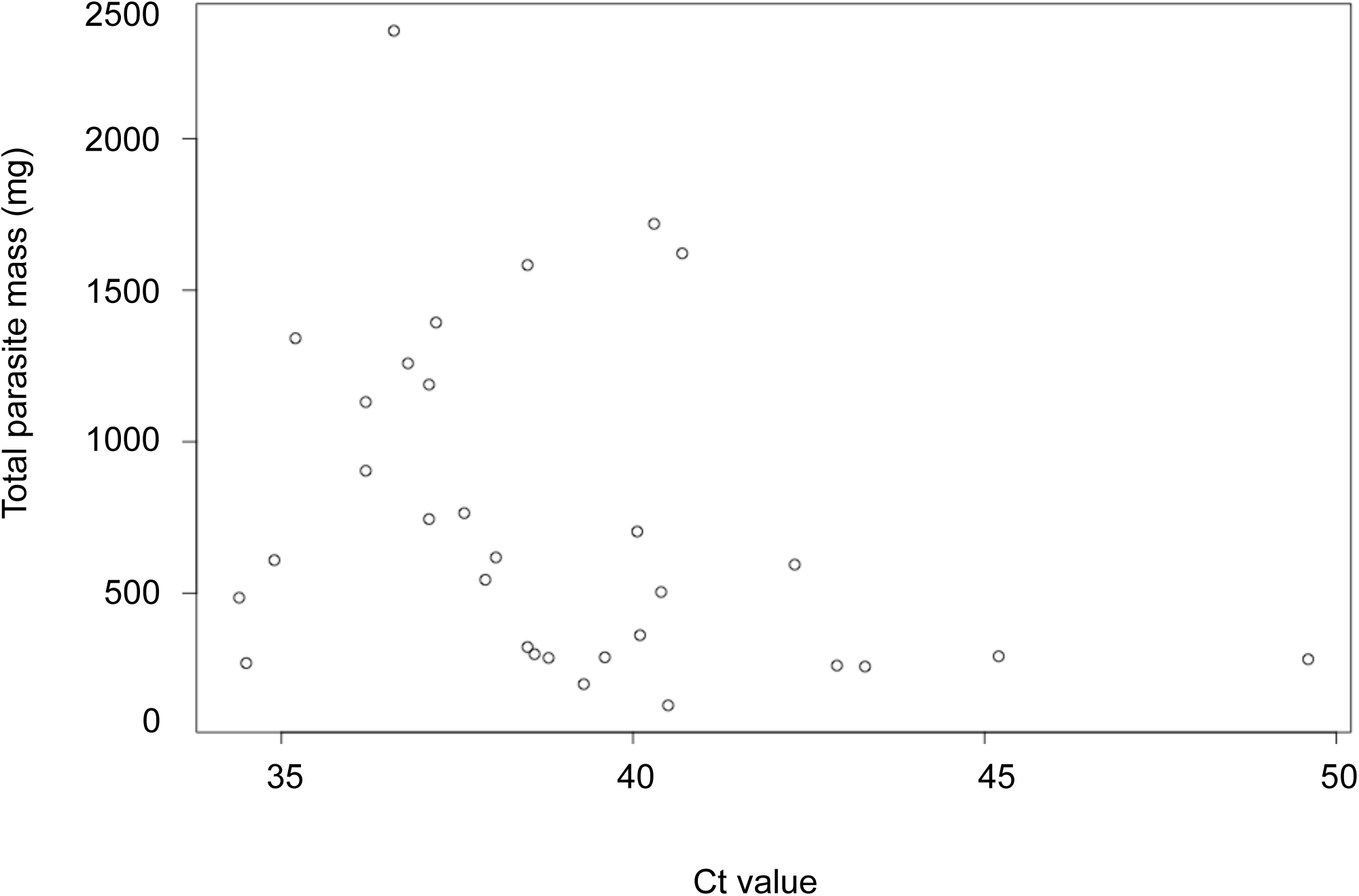
eDNA concentration covaries with total parasite mass. Low Ct values, which represent high levels of eDNA, are associated with a high parasite mass (Spearman correlation, rho = −0.41, p = 0.01, n= 31).

### Variations of ventilation rates between uninfected and infected sticklebacks

We measured the ventilation rates of 20 fish, 3 times over 3 consecutive days, and then determined that 13 of these were infected. The infected individuals harboured from 1 to 5 worms with a total mass ranging between 618 mg and 2356 mg (parasite index between 13.2 and 40.4). One infected individual died after the first ventilation measurement and we could use the videos for only the 2 first days for one of the non-infected fish. We had 3 ventilation measurements for all other individuals. Repeatability of ventilation across 3 days for a given fish was 0.41 (Lessells and Boag, 1987). As ventilation rates were repeatable over this time scale, we used ventilation rates measured on the first day in subsequent analysis, to reflect how this method would be used to quickly detect infection. Ventilation rates estimated over 30 seconds and reported in number of opercular movement per minute ranged between 99 and 146 opercular beats/min in uninfected fish and between 112 and 160 in *S. solidus* infected ones (Figure 3). The average number of ventilation per minute was 122 for uninfected fish and 134 for infected fish. Ventilation rates of uninfected fish were in the same range as found in previous studies of freshwater sticklebacks (Bell et al., 2010; Di Poi et al., 2016). We found that ventilation rates did not differ significantly between non-infected and infected individuals (two-sided t-test, t = −1.57, p = 0.15, n = 20, Figure 3). Ventilation was positively correlated to the parasite index, but this covariation was not significant (one-sided Spearman correlation, rho = 0.35, p = 0.07, n=20). There was a substantial overlap in ventilation rate between the two groups, which prevented the use of this trait to discriminate between *S. solidus* infected and healthy individuals. Therefore, the power of this simple measurement is low and can only be used as a complementary indicator of infection by *S. solidus* in its host.

**Figure 3.**
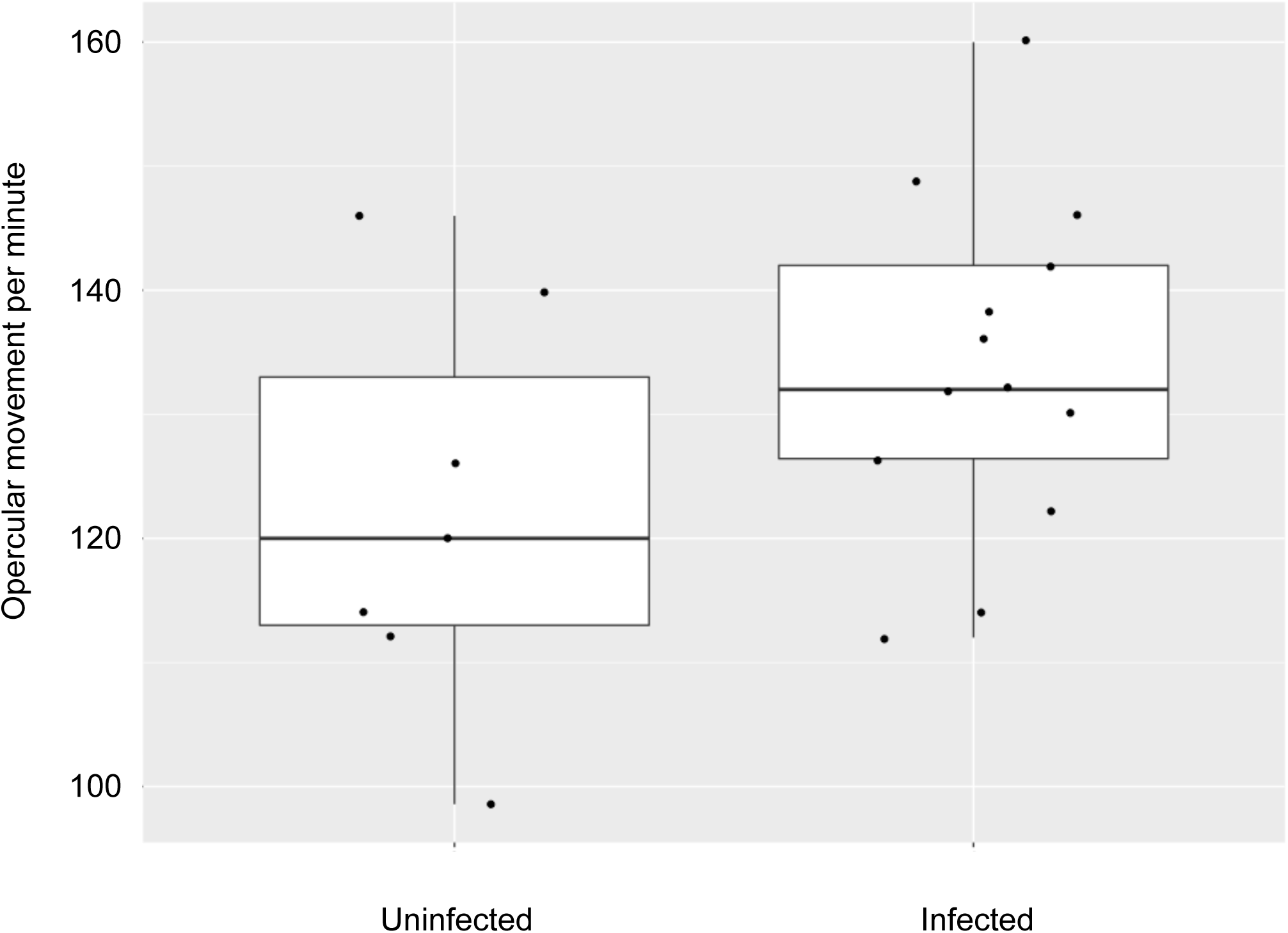
Ventilation rates did not differ significantly between un-infected and infected threespine sticklebacks. (two-sided t-test, t = −1.57, p = 0.15, n=20). Ventilation per minute ranged between 99 and 146 opercular movement per minute in un-infected fish (n=7) and between 112 and 160 in *S. solidus* infected fish (n=13). The lower and upper hinges correspond to the 25th and 75th percentiles and the vertical bars extend to the largest value no further than 1.5 times the interquartile range.

We expect the eDNA method presented here to find applications beyond the *S. solidus*-threespine stickleback system. Indeed, several fish species have parasites found in their abdominal cavity (Hoffman, 1999). To apply our method to these systems, the main challenge would be to design highly specific primers for the parasite of interest (Taylor et al., 2013). Our low cost method could be a simple alternative to morphological-based methods already used in these systems.

## Acknowledgments

The authors thank the personnel of the Laboratoire Régional des Sciences Aquatiques (LARSA) at Université Laval for their help with fish rearing. We thank AnaÏs Lacoursière-Roussel, Noémie Leduc, Alysse Perreault-Payette and Guillaume Côté for their advice concerning eDNA extraction. We thank Christian Landry and Leonard Foster for their comments during the development of this project. Anna Prudova from the Foster lab first suggested to us to sample the stickleback abdominal cavity fluids to study the *Schistocephalus* secretome, which led to this eDNA detection protocol.

## Author contributions

C.S.B. designed the study with input from N.A.-H. C.S.B. performed the eDNA-qPCR experiments and analyzed the data. N.A.-H. performed the ventilation rate measurements and did the statistical analyses. C.S.B. and N.A-H. wrote the manuscript.

## Ethics

This study was approved by the Comité de Protection des Animaux de l’Université Laval (CPAUL 2014069-3).

## Data availability

The complete dataset is available as a supplementary data file.

